# Selecting an effective amplitude threshold for neural spike detection

**DOI:** 10.1101/2022.01.25.477685

**Authors:** Zheng Zhang, Timothy G. Constandinou

**Affiliations:** Department of Electrical and Electronic Engineering, Imperial College London, South Kensington Campus, London SW7 2AZ. UK.; UK Dementia Research Institute (Care Research and Technology Centre) at Imperial College London and the University of Surrey

## Abstract

This paper assesses and challenges whether commonly used methods for defining amplitude thresholds for spike detection are optimal. This is achieved through empirical testing of single amplitude thresholds across multiple recordings of varying SNR levels. Our results suggest that the most widely used noise-statistics-driven threshold can suffer from parameter deviation in different noise levels. The spike-noise-driven threshold can be an ideal approach to set the threshold for spike detection, which suffers less from the parameter deviation and is robust to sub-optimal settings.

## I. Introduction

Brain Machine Interfaces are achieving significant new capabilities [1], [2] using neural spike waveform or spike timing in extracellular recordings. Spike detection is an essential step in extracting neural spikes from recordings. It not only distils the information for neural activity decoding but also reduces the data bandwidth for hundreds or even thousands times, making it possible for wireless transmission and achieving fully implant neural interfaces without percutaneous wires breaching the skin.

The spike detection performance is critical for preserving neural information and to avoid degradation of decoding accuracy. Thresholding is the most common way for spike detection and the values exceeding the threshold are regarded as spikes. An adaptive and robust threshold is then crucial facing the varying brain environment. There have been numerous algorithms proposed in the literature for defining the threshold. One approach is to use the computational algorithms [3], [4], for example, Short-time Fourier Transform, Wavelet Transform and template matching. There are also some algorithmic approaches, for example, a feedback-controlled threshold [5]. The most common approach is to set the threshold according to the signal statistics. The noise statistics are widely used to set a threshold. A hardware efficient estimation method is also proposed using a multiplier to set the mean/median/standard deviation/root-mean-square values as the threshold [6]. Others opt to set the threshold with a robust statistical estimation [7].

Setting the threshold as *T* = *αN*, where *N* is the noise statistics and *α* is a user-defined parameter, is a commonly used approach of setting the threshold [8]. This approach is especially preferred in on-implant implementation because of its simplicity [9]. However, the adaptiveness of such an approach is of concern. Some studies are seeking the optimal threshold for spike detection. In [10], it found that the noise mean value is the best threshold driving factor among other noise statistics. However, a single optimal multiplier that is suitable for varying SNR levels cannot be defined. In [5], it is assumed that the neural recording distribution consists of an exponent component (noise) and a power component (spikes). Here, a threshold is set at the intersection point of two estimated distributions. The distribution estimation can be resource-and-power-hunger in hardware, but the threshold driven jointly by noise and spike information can be heuristic. However, we have not seen anyone adopt the spike information to set the threshold computationally efficient or assess its suitability as a threshold driving factor.

In this paper, we investigate the performance of different multipliers based on noise and spike statistics across varying SNR levels. This allows us to assess the suitability to extracting spike information and how it compares to only using the noise level to set the threshold. Section II describes the dataset, performance metric and threshold settings. Section III shows the results we obtained from different threshold settings. Section IV discusses the findings from the results and analysis the challenges, and Sections V concludes this work.

## II. Methodology

### A. Synthetic dataset

In order to have more comprehensive control of the signal characteristics, we have generated a synthetic dataset in different noise levels for assessing the detection performance. The synthesising procedure is based on [8]. We use real recordings [11] from the motor cortex of monkeys sampled at 24,414 Hz to generate a synthetic dataset with ground truth. Synthetic recordings are consist of noise and spikes. The noise is truncated from the LFP removed and zero-centered real neural recordings in which periods spikes do not appear. The noise standard deviation (STD) is normalised to 1 and modified according to the desired SNR. The spikes are extracted from the real recordings using WaveClus [12] and there are 1,000 different templates with varying amplitudes for selection. The arrival of spikes can be simulated as a Poisson distribution with λ equal to the firing rate. By simulating multiple Poisson distributions of spike arrivals, multi-unit activities can be generated. One spike is randomly selected from the template spikes and chained with former spikes at desired arrival time. The gaps will be filled with zeros. After chaining all spikes from different cells, the spike amplitude will be normalised according to the spike peak mean values resulting in the unit average peak amplitude.

One synthetic recording is defined with three parameters: firing rate, number of cells and signal-to-noise ratio (SNR). The firing rate is the single-cell spike rate which determines the λ of each Poisson distribution, and the cell number determines the number of Poisson distributions be simulated. SNR is defined as the ratio of the mean value of spike peak amplitude and the STD of the noise.

### B. Threshold and spike detection

In order to find the optimal threshold derived from the signal statistics, we have assumed to have the perfect estimation of the noise and spike statistics. Three indicators: *μ_noise_*, *σ_noise_, μ_peak_*, which are (absolute) noise mean, noise STD and spike peak mean, are used to set the threshold:

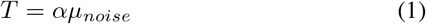

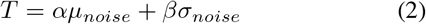

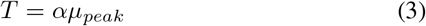

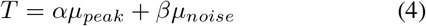

where *α* and *β* are user-defined parameters.

The detection occurs when the signal amplitude exceeds the threshold. As the spikes typical lasts for 20 timestamps, detection will be inactivated for 15 timestamps to avoid redetection. Detections falling within 10 samples around the ground truth are True Positives (TPs), others will are Flse Positives (FPs), undetected spikes are False Negatives (FNs).

### C. Evaluation metrics

To evaluate the performance of different threshold settings, we assess the detection accuracy (Acc), Span Ratio (SR) and Deviation-to-Span Ratio (DSR). Acc is formulated as:

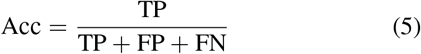

which describes the detection performance jointly considering sensitive and false detection rate.

*SR* describes the effect of an inaccurate set of the threshold parameters. The *Span* is defined the average multiplier difference in different noise levels that achieves the accepted accuracy. We prefer settings leading to large *Spans*, which means the multipliers values vs. Acc curve is flat, and it is less sensitive to the inaccurate setting of the threshold multipliers. However, the range of the multipliers in different settings can be different. To ensure a fair comparison among different settings, we take the ratio of *Span* at 0.8 Acc to the *Span* at 0.7 Acc as the metric *SR* to evaluate the robustness of inaccurate setting of the threshold within the same SNR level. More significant *SR* means better robustness. It can also be regarded as a measurement for the detection performance when the threshold is sub-optimal. The formula is given below:

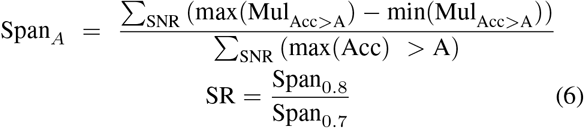

where *Mul* is multipliers, *max*(·) and *min*(·) are operators for max and min values, and A is the accepted accuracy level.

The *DSR* describes the deviation of the parameters across different SNRs for obtaining the best detection accuracy. *Dev* is the maximum difference for the multipliers that achieve the best detection accuracy higher than the accepted accuracy in different SNR levels. A small *Dev* means the best multiplier is less deviate from one SNR level to another. The adaptiveness of such a setting is therefore better in different SNR levels. Taking the ratio between the *Dev* and *Span* makes it possible to compare the settings with different parameter space. The *DSR* is defined as in Eq. 7 and we here assess the deviation at the accept accuracy level of 0.8, which gets involved in enough noise levels and fits the intuition of the minimum acceptable detection accuracy.

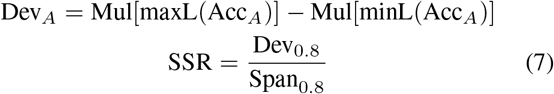

where *maxL*(·) and *minL*(·) are operations to find the lag of max and min values, and *Acc_A_* is the max detection accuracy in different SNR levels that are larger than A.

## III. Results

We have simulated 18 sets of recordings in which 2 cells each fires at 20 Hz. Their SNRs vary from 5 to 40 with a step of 2. Each set contains ten 4 s recordings. Threshold in each run is set with varying *α* and *β*. Spikes are detected according to the Section.II-B, and detection accuracy is averaged among 10 runs.

### A. *T* = *αμ_noise_*

We have swept *α* from 1 to 50, detection accuracy for different *α* and SNR is given in Fig. 1.A. Curves from top to bottom are cases SNR from high to low. The stars indicate the top settings that achieve the highest accuracy in different SNR. One can notice that there is a significant deviation for the optimal *α* from 15 to around 30 as the SNR increases. This indicates that the optimal threshold for spike detection does not increase linearly with the noise levels.

**Fig. 1.**
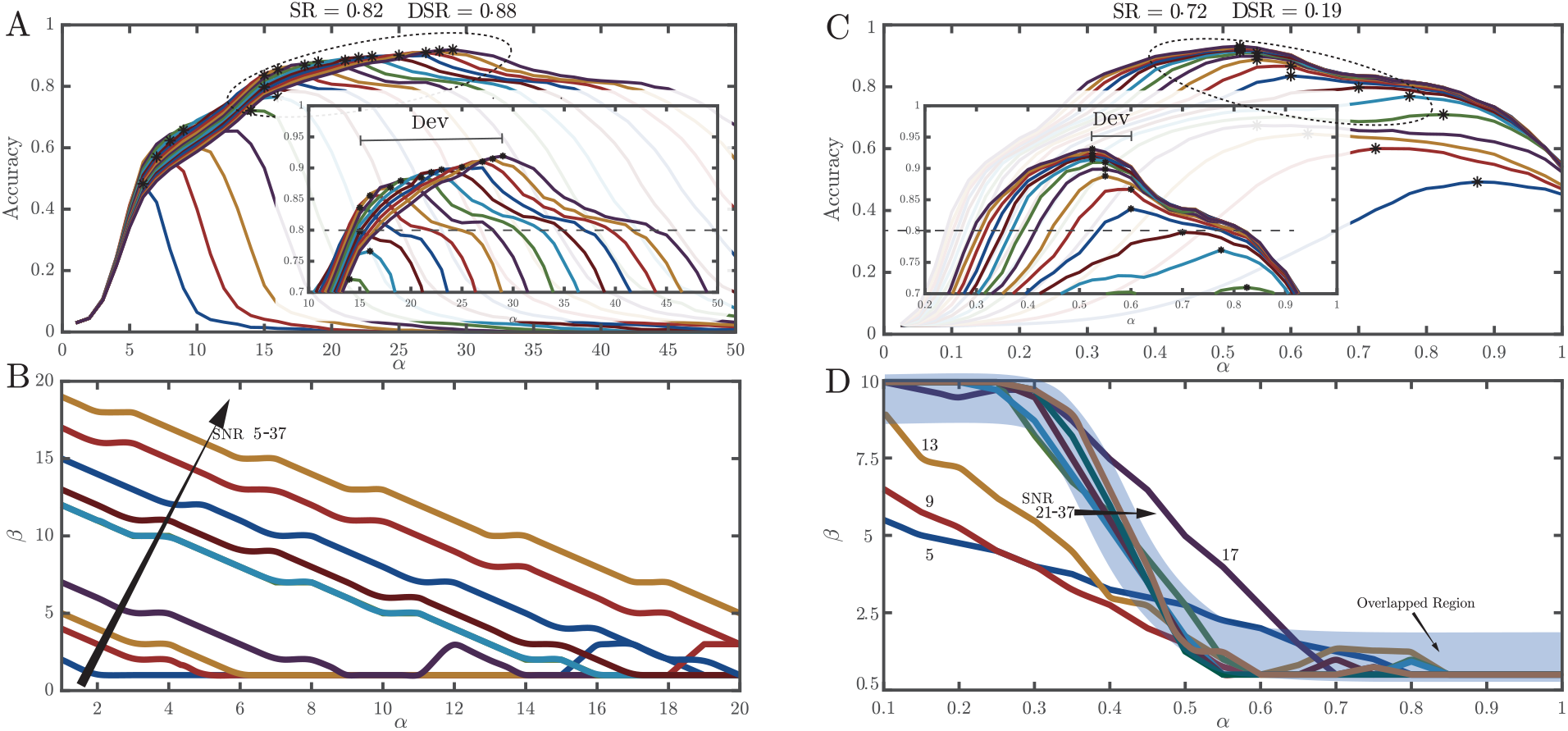
**A)** The detection accuracy using *T* = *αμ_noise_* for threshold with changing α and SNR. The curves from the bottom blue to the top purple are the SNR from 5 to 40 with a step of 2, and the stars indicate the top settings that achieve the highest accuracy in different SNR. (the same for C). **B)** The combination of *α* and *β* that achieves the best detection accuracy using *T* = *αμ_noiae_* + *βσ_noise_* for threshold in different SNR. The curves from bottom blue to top yellow are the SNR from 5 to 37 with a step of 4 (the same for D). **C)** The detection accuracy using *T* = *αμ_peak_* for threshold with changing *α* and SNR. **D)** The combination of *α* and *β* that achieves the best detection accuracy using *T* = *αμ_peak_* + *βμ_noise_* for threshold in different SNR. The overlapped region denotes the region of SNR levels and *α* – *β* combinations that suffers less from the parameter deviation for setting the optimal threshold.

### B. *T* = *αμ_noise_* + *βσ_noise_*

As the optimal threshold increases non-linearly with the noise mean, we tried to introduce the STD for thresholding. Both *α* and *β* are swept from 1 to 20. Results are shown in Fig. 1(B). Curves shown are the best combinations of the settings that achieve the highest accuracy in different SNR levels. The parameter deviation still exists in both the mean and STD sides as there is no overlapping among the settings in different SNRs. We therefore question if the noise statistic is a good or only indicator for setting the threshold.

### C. *T* = *αμ_spike_*

Intuitively, one can set a suitable threshold in the medway of spike peaks and noise ground, which means the spike peak level can be an indicator for setting the threshold. The *α* in Eq. 3 is set to varying from 0.1 to 2 with a step of 0.05. Results are shown in Fig. 1(C). DSR, which describes the parameter deviation, is reduced dramatically compared to Fig. 1(A), indicating that the spike guided threshold suffers much less than the noise-based threshold in different SNR levels. However, the SR is reduced, which means the threshold is more sensitive to the inaccurate set of the threshold. One reason is that the spike peak mean is high, and minor changes in the multiplier can significantly change the threshold level; another reason is that this threshold is noise invariant. When the noise is high, some large noises can exceed the threshold to increase the false detection.

### D. *T* = *αμ_spike_* + *βμ_noise_*

We can introduce noise awareness to the threshold by combining spike and noise statistics. With Eq. 4 in which *α* varies between 0.1 to 2 and *β* varies between 0.5 to 10, the top combination achieving the highest accuracy is shown in Fig.1(D). The deviation is much less than using Eq. 2 compared to Fig. 1(B). The accepted parameters are overlapped at the shaded blue region, which means the SNR in this region shares close optimal threshold settings. We selected three *α* values 0.25, 0.5, 0.6 and swept the *β*. The results are shown in Fig. 2. It can be observed that when *α* is 0.5, the DSR is minimal, and SR is also increased compared to Fig. 1(C). The parameter deviation can be increased when *α* deviates to 0.5 as DSR increases. However, the SR will still be maintained, which means the performance of different settings is consistent when the threshold is sub-optimal.

**Fig. 2.**
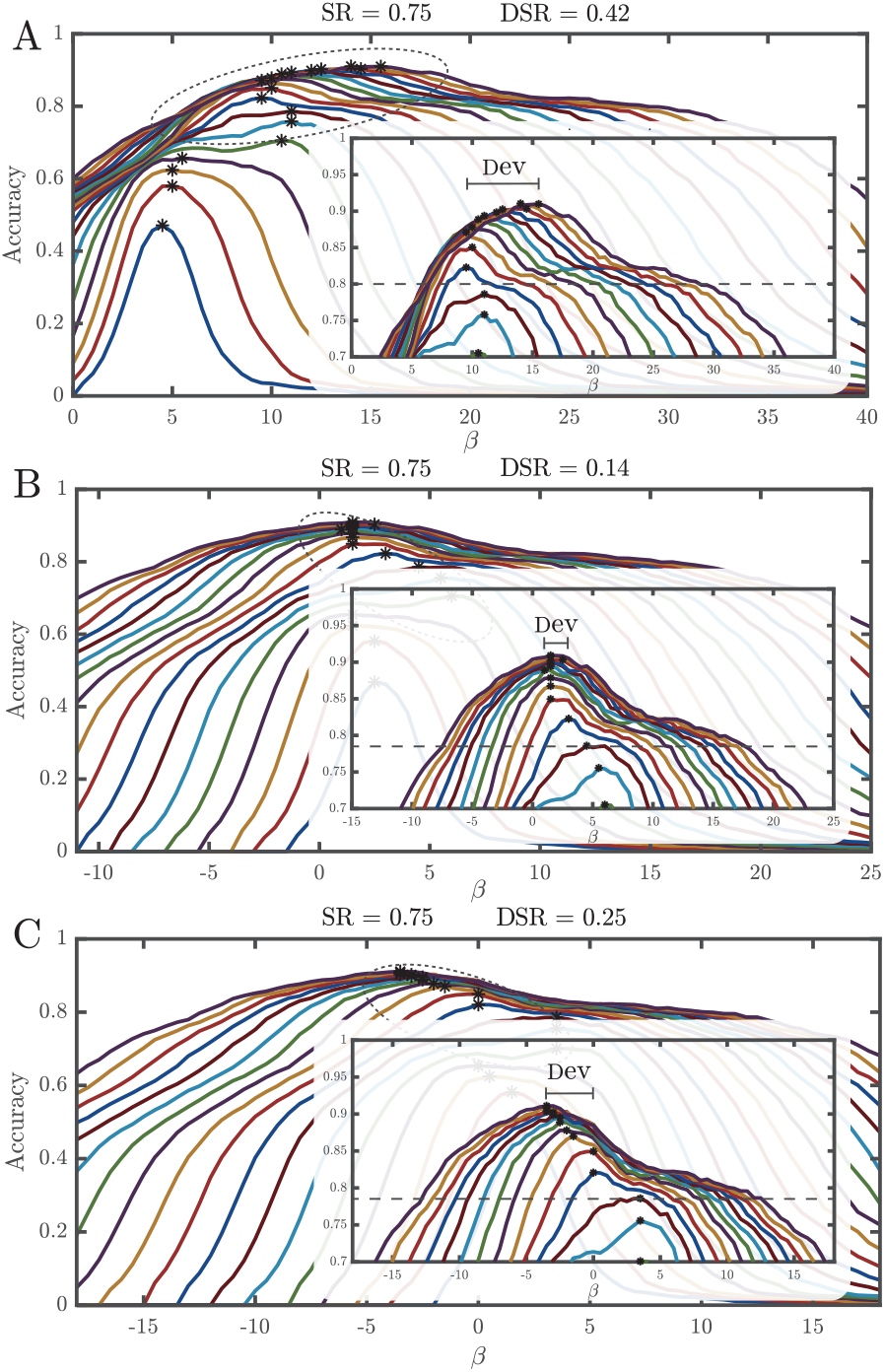
The detection accuracy using *T* = *αμ_peak_* + *βμ_noise_* for threshold with changing fix *α*, and varying β and SNR. The curves from the bottom blue to the top purple are different trials when SNR is increased from 5 to 40 with a step of 2. **A)** *α* = 0.25. **B)** *α* = 0.5,**C)** *α* = 0.6.

Such a joint spike-noise-based threshold utilises the peak values to set the coarse-grained baseline level of the threshold and uses the noise value to fine-tune the threshold. The coarse-grained baseline reduces the parameter deviation, and the fine-tuning provides noise awareness and increases the low SNR performance. Such a finding reveals the essence of the optimal thresholding and tells us we have forgotten an important factor - the spike peak in the past. Moreover, this also fits our intuition as the threshold should be SNR-driven.

### E. Application of the spike-noise based threshold to an independent dataset

According to Fig. 2(B), the suitable settings for *α* and *β* can be 0.5 and 1.5, respectively. In order to evaluate the generalisation of such a finding, we have applied these settings to a different dataset generated in [8], which has been widely used in multiple literature [6], [13]. The results are shown in the Table I. Compared to nearby settings, the maximum threshold is still peak at the selected settings, which mean there is no deviation between this dataset and the dataset we generated. Compared to other works, such a method achieves the highest detection accuracy and better robustness to different noise levels, referring to the mean and STD values of the Acc in different noise levels.

**TABLE I.** A COMPARISON OF DIFFERENT SPIKE DETECTION METHODS IN VARYING SNR

| Noise level | 0.05 | 0.1 | 0.15 | 0.2 | Mean | STD |
| --- | --- | --- | --- | --- | --- | --- |
| $\beta = 1$ | 0.98 | 0.99 | 0.96 | 0.88 | 0.953 | 0.05 |
| $\beta = 1.5$ | 0.99 | 0.99 | 0.97 | 0.92 | 0.968 | 0.031 |
| $\beta = 2$ | 0.99 | 0.99 | 0.96 | 0.92 | 0.967 | 0.031 |
| [6] | 0.89 | 0.97 | 0.92 | 0.89 | 0.918 | 0.037 |
| [8] | 0.92 | 0.95 | 0.95 | 0.85 | 0.918 | 0.047 |

## IV. Discussion

### A. Noise-driven threshold

The optimal threshold set only with noise statistics suffers severely from the changing noise levels. The parameter deviation lowers the model adaptiveness. The reason is that noise statistics is not the only dependent factor to the optimal threshold. The best setting for one SNR can overfit such noise level and no longer works with the noise level changes.

### B. Spike-driven threshold

Using spike peak in guidance of set the threshold can overcome parameter deviation. A suitable parameter that generalises well in different noise levels can be found. However, as the spike values are more significant than the noise, the threshold becomes more sensitive to the change of multiplier values. When the threshold is sub-optimal, the detection performance can be affected.

### C. Spike-noise-driven threshold

Jointly using the spike and noise statistics can trade-off between parameter deviation and sub-optimal threshold degradation. Using spike values can effectively reduce parameter deviation, and using noise values reduces the effect of sub-optimal threshold on degrading detection performance.

### D. The concern on spike peak estimation

Such an approach requires robust estimation on both noise and spikes. There are plenty of researches focusing on noise estimation but few studies are working on the spike peak estimation. Estimating the spike peak can be challenging as we have no prior knowledge of the peak amplitudes before detection. The threshold will be updated according to the local environment in real-time adaptive spike detection. Without a robust estimation of the spike peak amplitude, the false detection of the spikes could lower the estimated spikes peak amplitude and even lower the threshold leading to more false detections. Such positive feedback could eventually crush the spike detection algorithm. A Kalman filter or PID control can potential be used for the spike peak estimation.

### E. The concern on the optimal threshold

The optimal threshold setting we found in this paper is *T* = 0.5*μ_peak_* + 1.5*μ_noise_*. However, it still needs the verification of its suitability on the real signal recordings, which can be challenging as there is no ground truth label. There is also the need for a proof of whether 0.5 and 1.5 are good estimation for spike detection using Eq. 4.

## V. Conclusions

In this paper, we have explored the factors determining the optimal threshold for spike detection. We have found that an optimal threshold that generalises in the different SNRs should be derived from both spike peak amplitudes and noises. We also point out that the peak amplitude estimation can be challenging. Without a robust estimation, the optimal threshold can hardly be achieved. That leads us to our future works on robust spike peak estimation and other thresholding techniques independent from the noise or spikes.

